# Micropropagation and Assessment of Genetic Homogeneity in *Dendrobium moniliforme* (L.) Sw. using Molecular Markers

**DOI:** 10.1101/2025.08.31.673335

**Authors:** Manisha Ghimire, Mahendra Thapa, Pusp Raj Joshi, Mukti Ram Paudel, Krishna Kumar Pant, Pritam Gurung, Sven H. Wagner, Bijaya Pant

## Abstract

*Dendrobium moniliforme* (L.) Sw., is a valuable medicinal orchid widely used as tonic. This study aimed to develop an effective protocol for micropropagation of *D. moniliforme* and to compare the genetic similarities of the micro-propagated plants to the wild plants using Random Amplified Polymorphic DNA (RAPD) and Inter Simple Sequence Repeats (ISSR) molecular markers.

Healthy seeds were cultured on Murashige-Skoog (MS) medium, supplemented with various concentrations of auxins, cytokinins and coconut water for shoot and root development. Genetic homogeneity was assessed using ten primers of each RAPD and ISSR marker.

MS media fortified with 10% coconut water was efficient for seed germination and protocorm development. Maximum number of shoots were produced on half strength MS (HMS) medium supplemented with 0.25 mg/L NAA and 10% coconut water (CW) i.e 22 shoots. FMS supplemented with 0.5 mg/L indole butyric acid (IBA) was found to be best as plant hormone for root induction, as it induced ten roots/shoot. RAPD and ISSR amplified a total of 15 and 5 homogeneous DNA fragments. This protocol demonstrates a reliable and efficient method for large-scale propagation and conservation of *D. moniliforme*, supporting its commercial and medicinal value.

**Highlights:** *Dendrobium moniliforme* was micro-propagated from seeds using MS medium. RAPD and ISSR markers confirmed its genetic stability. The study supports large-scale true-to-type propagation and conservation of these valuable medicinal orchids.

## 1. Introduction

*Dendrobium moniliforme* (L.) Sw. is widely distributed across the subtropical and temperate regions of South and Southeast Asia (Pant et al., 2020). The flower has pale yellow to whitish petals with purple or red spots (Figure 1A and 1B). It is a valuable orchid species known for its medicinal and horticultural significance.

**Figure 1.**
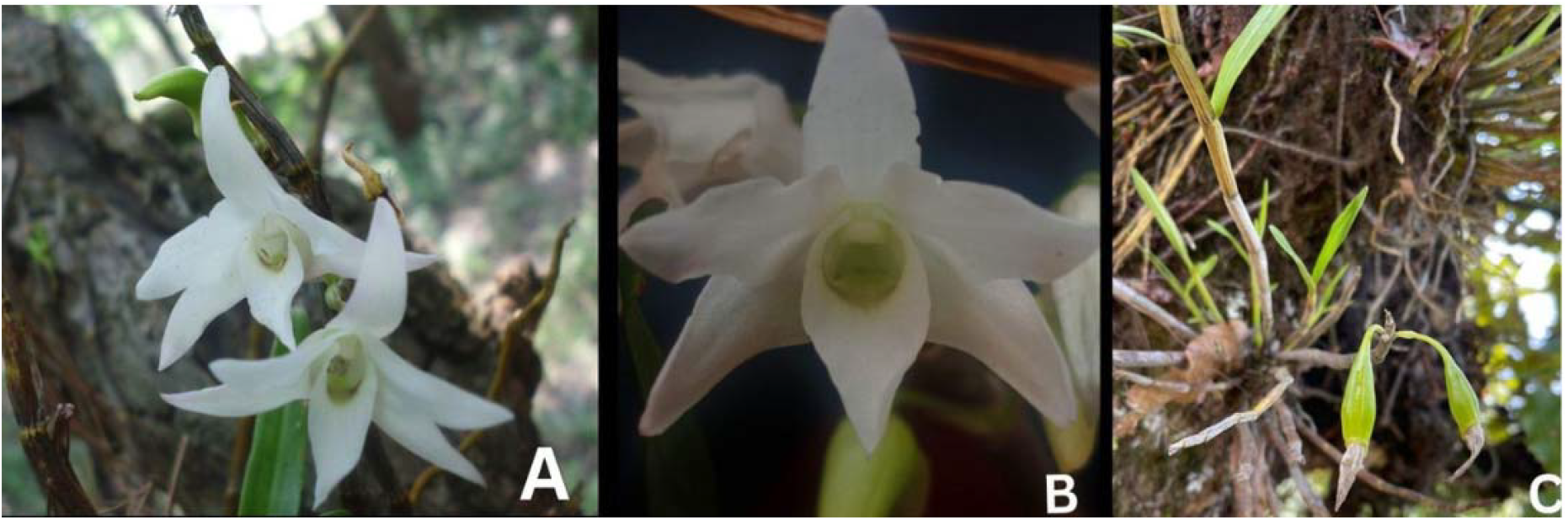
Plants of *Dendrobium moniliforme* (L.) Sw.; A-C: *D.moniliforme* in natural habitat; B. Flower of *D. moniliforme* showing white and the labellum or the lip is green with brownish-purple spots; C: Pod of *D. moniliforme*.

It has been utilized as a peculiar ingredient in a number of traditional Chinese Medicines for the formulation of remedies to cure ailments related to the eye, as antipyretics, and tonics (Wu et al. 2014). For instance, a well-known traditional Chinese medicine, and known as- “Tongpi Fengdou,” is produced from the stems of *Dendrobium moniliforme* (Hu, 1970; Zhang et al. 2011). The plant is also used to cure several pulmonary and gastro-intestinal issues by different ethnic communities of the Indian Subcontinent (Wang et al. 2021). The specific bioactive compounds in *D. moniliforme* are polyphenolic dendrocandins, denbinobin and squaric acid derivative: moniliformin. These compounds are known for several pharmacological effects including anti-osteoporosis (Baek et al. 2016), antioxidant and anticancer properties (Paudel et al. 2018; Pant et al. 2021). Orchids are among the most valuable and widely traded flowers in the global floriculture industry (Hsing et al. 2016; Liang et al. 2020). These values are significantly increasing their global demand leading to their excessive collection. Orchids are habitat-specific and require specialized ecological interactions, such as specific pollinators or mycorrhiza, to maintain their life cycle. Moreover, natural seed germination is extremely limited, occurring at a rate of less than 3% due to its dependence on mycorrhizal fungi (Vyas et al. 2009). These factors make them vulnerable and susceptible to extinction. As a result, the whole family is listed under Appendix-II CITIES, and the majority of species are declared in the IUCN Red Lists category (IUCN, 2020). The cultivation and production of these orchids are globally increasing (Yuan et al. 2021). Conventional approaches are less effective since the adequate vegetative propagation or propagation via seeds is practically impossible meeting the global demand for these flowers. Micropropagation could be a promising option to address this issue because it provides a reliable means for rapid, large-scale production of plantlets and allows continuous propagation throughout the year (Pant 2013; 2014). The optimization process involves altering several variables, including the culture conditions (temperature, light and humidity), the explant source selection, the growth medium and hormone selection, and the sterilization procedures (Nanaware et al. 2024).

However, micropropagation is prone to several alterations in plantlets at the genetic level. Most explants subjected to micropropagation exhibited somaclonal variation, a phenomenon where genetically identical plant clones display different physical traits after regeneration. This variation becomes more pronounced with prolonged cell culture propagation and repeated sub-culturing, which can stress the explants and micropropagated plantlets (Ferriera et al. 2023). Among the variety of factors of clonal variations, the source of explant and types of plant regulators are the most significant ones (Devi et al. 2014).

The tools to explore the genetic stability of micro-propagated plantlets have been extensively studied. These commonly include cytological, microscopic, and molecular techniques. Molecular markers offer robust approaches for precise and quick identification of somaclonal variations (Cloutier & Landry, 1994), of which RAPD and ISSR are commonly used. RAPD and ISSR are low-cost, easy to use, and do not require any previous DNA sequence information. RAPD which relies on the amplification of DNA, uses short, arbitrary primers (usually 10-20 base pairs in length) that can anneal to random genomic DNA sequences in a simple, easy and cost-effective manner (Williams et al. 1990). ISSR uses a single primer, 16–20 base pair in length made up of repetitive bases, which are placed at the 3′ or 5′ end of 2-4 random oligonucleotides (Javan and Heidari, 2012). The somaclonal variation in plant tissue culture-induced plantlets is studied in some species of *Dendrobium*, such as *D. moniliforme* using an Amplified Fragment Length Polymorphism (AFLP) marker (Ye et al. 2015), and *D. transparens* using RAPD and ISSR markers (Joshi et al. 2023). This research aims to support the development of a comprehensive protocol for the conservation and large-scale production of genetically stable *Dendrobium moniliforme* plantlets to meet existing medicinal and industrial demands. Additionally, there have been very few studies accomplished on the genetic homogeneity of *in vitro* plantlets, particularly in *D. moniliforme*. This study focuses on regenerating *D. moniliforme* plantlets using MS medium with various plant growth regulators (PGRs), acclimatizing the regenerated plants, and verifying their genetic similarity to the mother plant.

## 2. Materials and methods

### 2.1 Mass propagation of *D. moniliforme*

#### 2.1.1 Sample collection

*D. moniliforme* was collected from the Phulchowki of Lalitpur District of Central Nepal in March 2023. The collected plants were processed for herbarium preparation to ensure accurate identification (Lawrence, 2017). The voucher specimen of herbarium (voucher no.M02) was deposited at the Tribhuvan University Central Herbarium (TUCH), Kirtipur, Nepal. The pods of this orchid (Figure 1C) were used for tissue culture.

#### 2.1.2 Media preparation

Different strengths of MS media: full strength (FMS), half strength (HMS), and quarter strength (QMS) (Murashige and Skoog, 1962) were used applied. These media were also supplemented with auxins (IAA, IBA, and NAA), cytokinins (BAP and Kinetin), and coconut water (CW). The pH of all media was adjusted to 5.8 using either 0.1N NaOH or 0.1N HCl before autoclaving, and then 0.8% agar-agar powder (ThermoFisher Scientific India) was added. The prepared media were poured into either jars or tubes. Approximately 50ml of media was used per jar, and 20ml per tube. The mouths of the jars or tubes were covered with transparent lids or aluminium foil and autoclaved for 20 minutes at 121°C and at 15 pounds per square inch (psi) above normal atmospheric pressure. The sterilized media were then kept at room temperature for one week before inoculation.

#### 2.1.3 Surface sterilization

The mature pods of *D.moniliforme* were thoroughly washed with one to two drops of Tween-20 under running tap water for 30-40 minutes. They were then transferred to a laminar flow cabinet for surface sterilization, which involved dipping them in 0.1% mercuric chloride solution for 15 minutes, followed by 70% ethanol for 2 min. Afterwards, the pods were rinsed three times with sterile distilled water. Finally, they were placed on a sterile filter paper in a sterile petri dish to remove excess moisture.

#### 2.1.4 Inoculation and maintenance of culture

The immature pods were longitudinally cut with a sterile blade and the powdery seeds were inoculated on different media combinations-FMS, HMS, and QMS alone and in combination with phytohormones-0.5 mg/l BAP and 0.1 mg/l NAA and 10%CW separately. The inoculated explants were transferred to a culture room at a temperature of 25 ±3 °C, 16 hours of illumination, and a light intensity of 73.17 to 97.56 micromole per square meter per second (µM/m^2^/s) using cool white fluorescent tube lights.

#### 2.1.5 Shoot multiplication

For shoot multiplication, protocorms were inoculated on FMS medium and HMS medium, respectively, alone or in combination with NAA and BAP, at a concentration range from 0.25 to 2.0 mg/L different combinations for shoot development. The efficacy of using coconut water for the shoot multiplication is also studied at different concentrations (10, 15, and 20%).

#### 2.1.6 Root induction

The propagated shoots were subsequently grown in MS media supplemented with varying concentrations (0.25 to 2 mg/L) of auxins (IAA, IBA, and NAA), resulting in a total of 12 different root induction media. Root development was measured in terms of root length and the number of new roots formed. The data were collected after the fourth week of inoculation. Morphological changes were recorded based on visual observations.

#### 2.1.7 Acclimatization

Plantlets above 3-5 cm in height with well-developed roots were selected for acclimatization. Jar containing plantlets were left in the greenhouse for two weeks for hardening. After two weeks the plantlets were gently extracted from culture vessels using sterile forceps to avoid root damage and rinsed in sterile water to remove residual agar from the root surfaces and finally transplanted into earthen pots containing a different ratio of potting mixture like cocopeat: moss at a 2:1 ratio, pine bark moss□at 2:1; cocopeat: moss: pinebark at -□3:1:1. The potted plantlets were maintained in a shade house under controlled humidity (70–80%) and indirect light, with ambient temperatures ranging from 26□°C to 27□°C.

### 2.2 Genetic stability study of *in vitro* developed plants compared with mother plants

#### 2.2.1 Selection of *in vitro* samples for DNA isolation

Altogether five different samples grown at different conditions were selected for DNA extraction. In vivo mother plant, in vitro plantlets grown on 0.25 mg/L NAA with 10% CW, 0.5 mg/L IBA, protocorm grown on basal medium, and acclimatized plantlets were selected for the genetic fidelity studies. The genetic profiling experiment was performed in Annapurna Research Center, Maitighar, Kathmandu, Nepal.

#### 2.2.2 Extraction of DNA by the CTAB method

Genomic DNA of *D. moniliforme* was isolated from both *in vitro* and *in vivo* samples using the CTAB method (Doyle, 1991), with a few changes. From each sample, 0.2 g of leaves, protocorms, was taken and crushed with liquid nitrogen using a mortar and pestle to make a fine paste. 500 µl of CTAB buffer (2 % CTAB, 0.5 mM EDTA, 5 mM NaCl, 1 mM Tris-HCl, pH 8.0, 0.2% -mercaptoethanol) was added to it. It was then poured into a clean, sterilized, 2 ml microcentrifuge tube. The samples were incubated for 45 minutes at 60 °C in a circulating water bath. After incubation, samples were centrifuged for five minutes at 10,000 rpm to settle the cell debris. Supernatants were then transferred to a clean, sterile microcentrifuge tube. An equal volume of chloroform:isoamyl alcohol was added, and the mixture was carefully mixed by inversion some times (5-8 min). It was then centrifuged for 10 minutes at 12,000 rpm. Supernatant was shifted to a micro-centrifuge tube, and an equal volume of ice-cold propanol was added and stored at -20 °C for an hour. DNA was centrifuged at 12,000 rpm for five minutes to form a pellet. 70% ethanol was added, centrifuged at 10,000 rpm for five minutes, and left to dehydrate for 15 minutes. Agarose gel electrophoresis was performed, and the quality of DNA was evaluated using a fluorometer (Thermo Fisher Scientific, USA). The concentration of extracted DNA was adjusted to 30ng and stored at -20 °C, and UV spectrophotometry was used to assess the amount and quality of extracted DNA.

#### 2.2.3 DNA amplification and RAPD, ISSR analysis

A total of ten RAPD and ISSR primers were used for the amplification of template DNA. Primers with a clear and reproducible banding pattern were taken for further processing. The amplification was done by using multiplex PCR. PCR reactions for RAPD and ISSR assays were performed in a 15 μl reaction volume. It includes 5.5 μl of nuclease-free water, 6.5μl of 2X PCR master mix (GeneDireX, Taiwan), 1 μl of primers (Macrogen, Korea) and 2 μl of DNA template. The cycling conditions were optimized using a PCR thermal cycle (ProFlex PCR, Thermo Fisher Scientific, USA) following Cerasela et al, 2011. The PCR cycling condition was programmed at 92 °C for 5 min, followed by 45 cycles of denaturing at 92 °C for 1 min, annealing at 37-40°C for 1 min, extension at 72°Cfor 1 min and final extension at 72°C for 5 min for RAPD and ISSR, initial denaturation 95°C for 5 min, followed by 35 cycles of denaturation at 94°C for 1 min, annealing of 45-50°C for 1 min, extension at 72 °C for 2 min, and then a final extension at 72 °C for 7 min.

To test the utility of the primers, PCR products were detected on 1.5% agarose gels with 1 μg/mL ethidium bromide in 1X TBE (Tris base, boric acid, and EDTA) buffer with a constant voltage of 70 V for 1.5 hr. The size of the amplification product was determined by comparison with the 100 bp DNA ladder marker (Thermo Fisher Scientific, USA). The clear and reproducible bands were considered for counting alleles.

### 2.3 Experimental design and statistical analysis

The experiment was conducted in a laboratory setting and repeated at least three times to ensure reproducibility. Visual observations were made every week. After four weeks of induction, protocorm formation and the number per explant were noted. Shoot development was assessed four weeks after shoot induction, and root formation was evaluated 25 days after root initiation. Data are presented as mean ±SD and analyzed using one-way analysis of variance (ANOVA) followed by Duncan’s multiple range test (p< 0.05) using SPSS (version 25).

For the genetic homogeneity test, RAPD and ISSR banding patterns were manually evaluated. Clear and reproducible bands were selected, and alleles were scored based on their presence (1) or absence (0) in each amplified DNA fragment. A binary matrix was created to compile the presence-absence data. Allele sizes were estimated using DNA ladder markers. Banding patterns from different markers were analyzed and compared individually.

## 3. Results

### 3.1 In vitro seed germination

Among all the tested combinations, full-strength MS medium fortified with 10% CW was found to be effective for seed germination and protocorm formation as it involved five weeks and approximately six weeks for the process respectively. This combination showed the shortest time required for the process. The media combination with phytohormones was found to be least effective. QMS added with 0.5 BAP and 0.1 NAA required a maximum of weeks, i.e. eight weeks for germination and eleven weeks for protocorm formation. The relative strength of MS media also played a role in the process. FMS, HMS, and QMS required approximately six, seven and seven weeks for seed germination and seven, seven and ten weeks, respectively for protocorm formation. Comparatively, 10%CW was positively significant for the consistently promoted faster protocorm development across all the media.

### 3.2 Shoot multiplication and elongation

#### 3.2.1 Role of phytohormones

The visible shoot initiation and development were noticeable only after three weeks of inoculation. Among, all the treatments, low concentration of NAA fortified with coconut water was found to be superior for shoot multiplication. HMS supplemented with 0.25 mg/L NAA+10% CW resulted in the highest number of shoots ie. 22 shoots/explant (Table 1). Additionally, NAA combined with 10% CW was effective than NAA alone or BAP. Among the BAP-subjected medium, half-strength MS supplemented with one BAP produced 18 shoots/ explant. HMS with 0.25 NAA produced 12 shoots/explant (Figure 2). The synergism between coconut water and BAP was found to be negative. BAP combined with 10% CW produced 11 fewer shoots than BAP alone. The strength of MS medium was found to be significant alone for the shoot multiplication. HMS produced 13 shoots/ explant, which is 3 shoots more than in FMS.

**Table 1.** Seed germination and protocorm formation in weeks of *D. moniliforme* on the different media combinations.

| Media Combinations<br>(PGRs in mg/L) | Seed Germination in<br>Weeks (Mean ± SE) | Protocorm Formation in<br>Weeks (Mean ± SE) |
| --- | --- | --- |
| FMS | 5.66±0.33<sup>ab</sup> | 7.0±0<sup>b</sup> |
| FMS+10% CW | 5.0±0<sup>a</sup> | 6.33±0.33<sup>a</sup> |
| FMS+0.25BAP+0.1NAA | 6.66±0.33<sup>cd</sup> | 9.0±0<sup>c</sup> |
| FMS+0.5BAP+0.1NAA | 7.0±0<sup>cd</sup> | 8.66±0.33<sup>c</sup> |
| HMS | 6.66±0.33<sup>cd</sup> | 7±0<sup>b</sup> |
| HMA+10% CW | 6.33±0.33<sup>bc</sup> | 7±0<sup>bc</sup> |
| HMS+0.25BAP+0.1NAA | 7.33±0.33<sup>de</sup> | 10±0<sup>d</sup> |
| HMS+0.5BAP+0.1NAA | 8.0±0<sup>e</sup> | 10±0<sup>d</sup> |
| QMS | 7.0±0<sup>cd</sup> | 10±0<sup>d</sup> |
| QMS+10% CW | 7.0±0<sup>cd</sup> | 9±0<sup>c</sup> |
| QMS+0.25BAP+0.1NAA | 6.33±0.33<sup>bc</sup> | 11±0<sup>e</sup> |
| QMS+0.5BAP+0.1NAA | 8.0±0<sup>e</sup> | 11±0<sup>e</sup> |

<sup>a-e</sup> Values with different superscripts are significantly different at $p \leq 0.05$ using Duncan's Multiple range tests (n=36).

**Figure 2.**
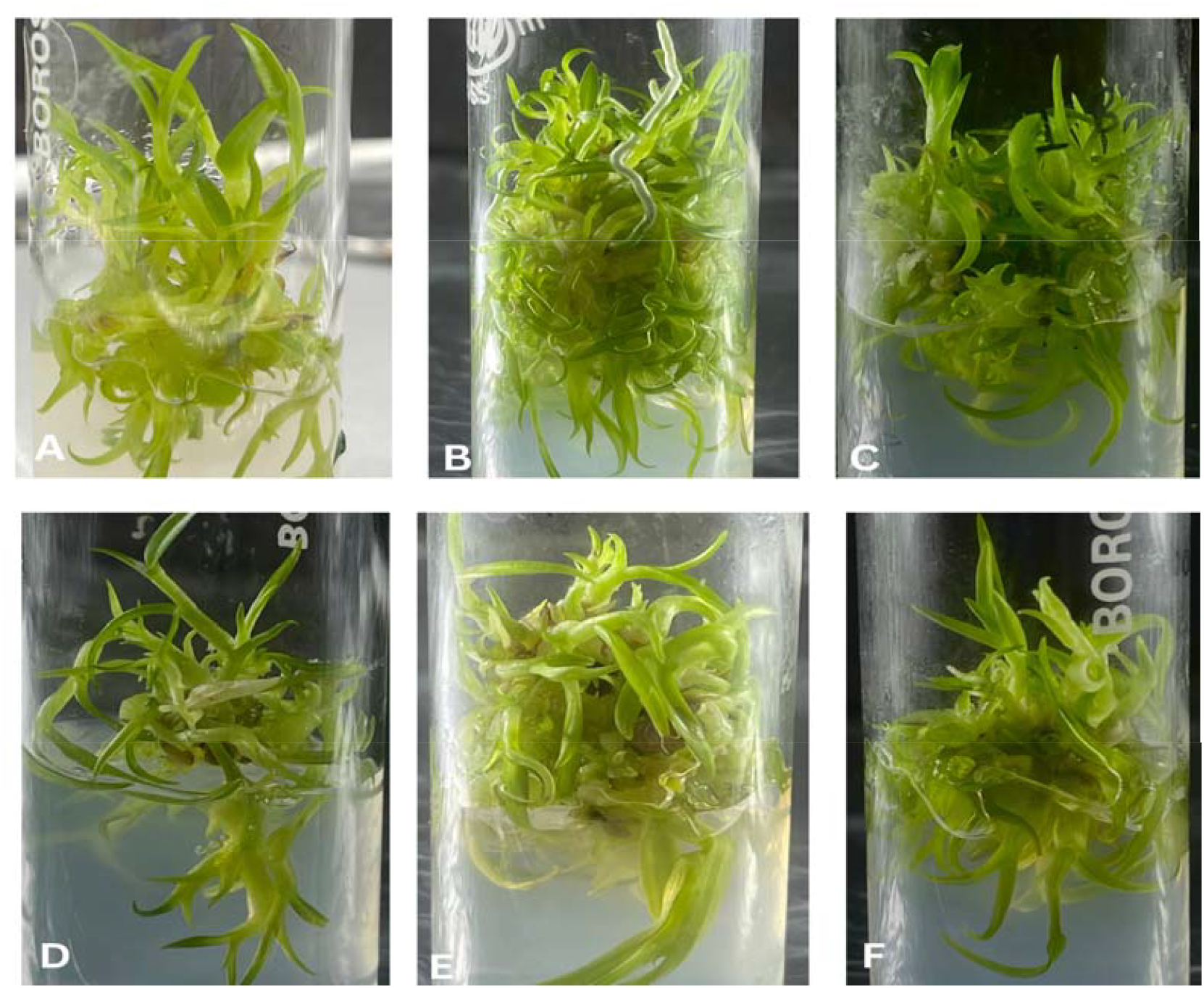
Effect of different concentrations of growth hormones and coconut water for shoot development A: HMS+0.25 mg/L NAA; B: HMS+ 0.25 mg/L NAA +10% CW; C: HMS+ 1 mg/L BAP; D: FMS; E: HMS+ 0.25 mg/L BAP; F: HMS

Regarding shoot elongation, maximum shoot elongation was found to be on HMS+ 0.25 NAA+10% CW as it elongated shoots by 2.16 cm (Table 2), followed by HMS+0.25BAP+10%CW (2.00±0.05). Additionally, NAA combined with 10% CW was effective than NAA alone or BAP. Least shoot elongation was shown by FMS with 10% CW, i.e 0.36 cm. The concentration of NAA was found to be crucial for both shoot elongation and new shoot formation when supplemented with coconut water.

**Table 2.**
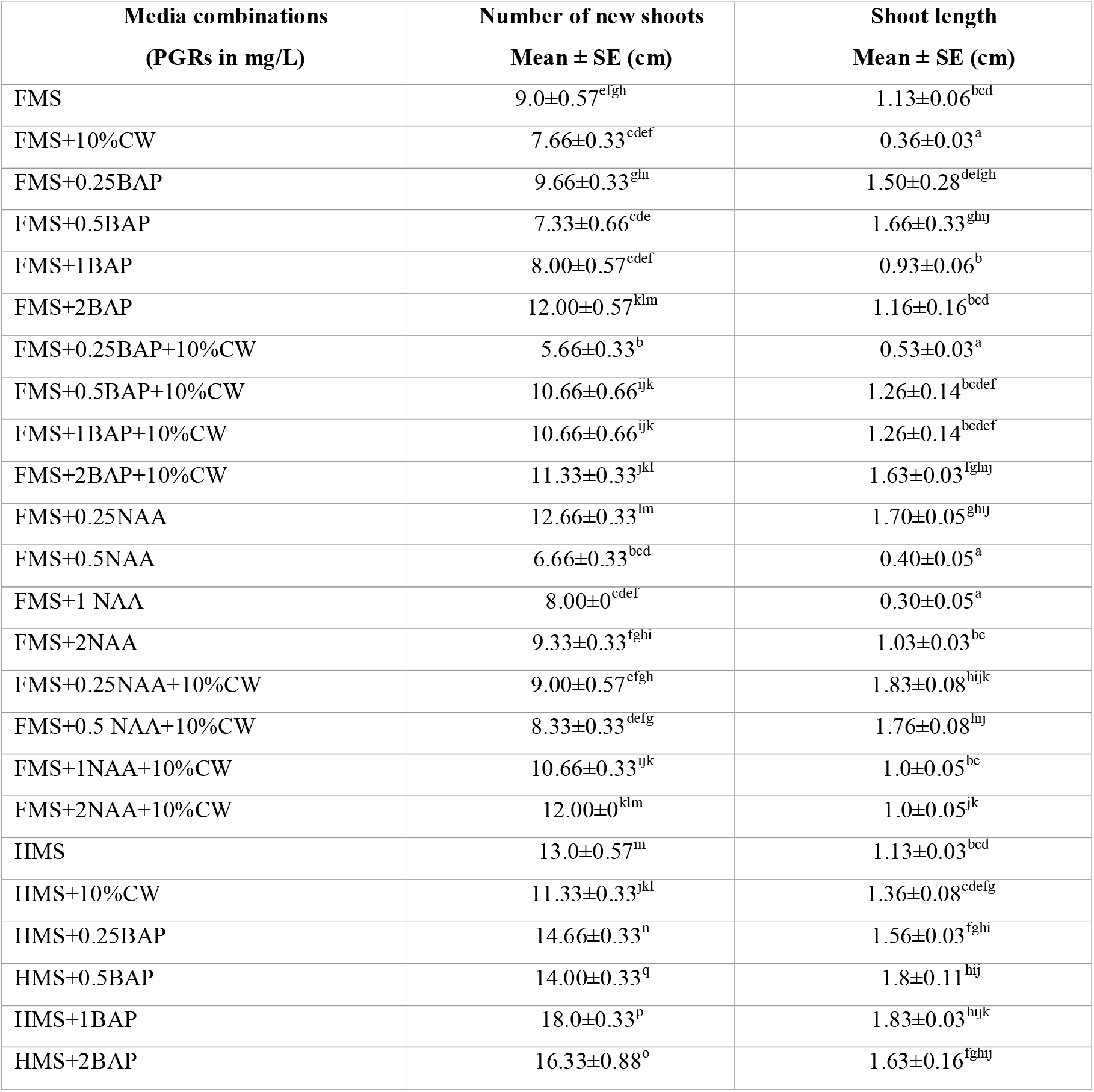
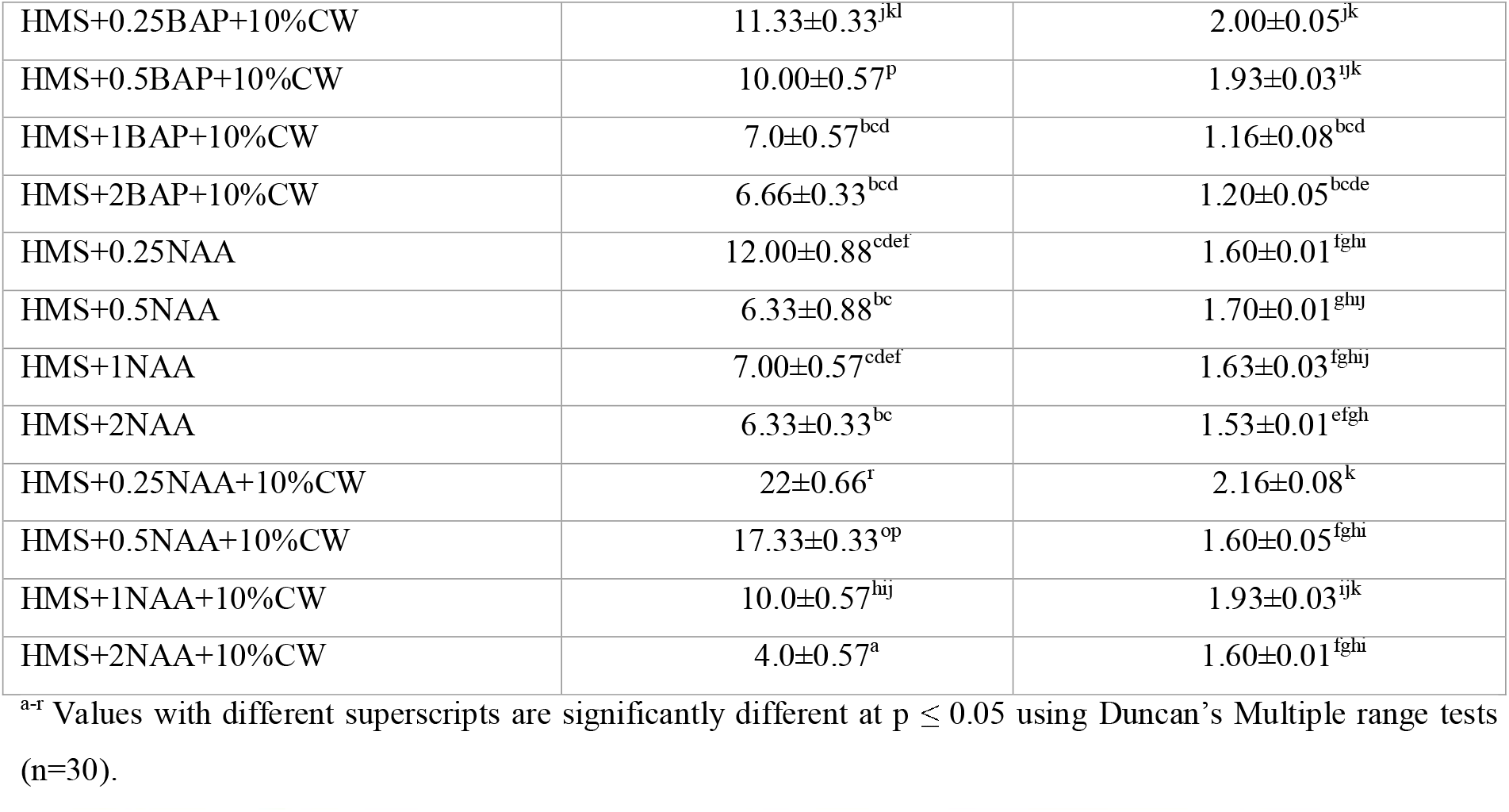
Shoot multiplication and elongation of *D. moniliforme* on the different media combinations.

#### 3.2.2 Role of different concentration of coconut water in shoot development

*In vitro* growth of shoots showed a remarkable response with the use of coconut water in terms of shoot multiplication and had minor effect on shoot elongation. Three different concentrations of coconut water 10%, 15% and 20% were tested in half-strength MS media. The tested coconut water concentration from 10-20% showed a similar gradual decreasing pattern for the new shoot formation under both HMS and FMS medium. Coconut water at the concentration of 10% produced maximum shoots on either FMS or HMS medium. HMS was found to be superior to FMS for the new shoot formation. HMS+10%CW produced an average of 15 shoots while FMS+10%CW produced around 8 shoots. Least shoot development on the coconut water-mediated medium was observed on FMS+20%CW. It produced an average of three shoots. This result is followed by FMS+15% CW with an average of five shoots. Concentrations beyond 10% CW showed decreased shoot growth Regarding shoot length, among different concentration of CW on both FMS and HMS, remains relatively constant with only slight variations.

### 3.3 In vitro root development

Selected auxins at different concentrations had a diverse effect on root development for several new roots and their elongation. Full strength of MS medium (FMS) with 0.5 mg/L IBA had best for the growth of new root. It produced an average of 10 roots/shoot which is followed by FMS+1.5 mg/L IBA where 8 roots/shoot were observed. The least number of roots (around two) was developed in a shoot on the FMS+2 mg/L NAA followed by FMS+0.5 mg/L IAA where around 3 roots were developed in a shoot. Among the tested auxins, IBA was significant for increasing root number (Figure 3 and 4).

**Figure 3.**
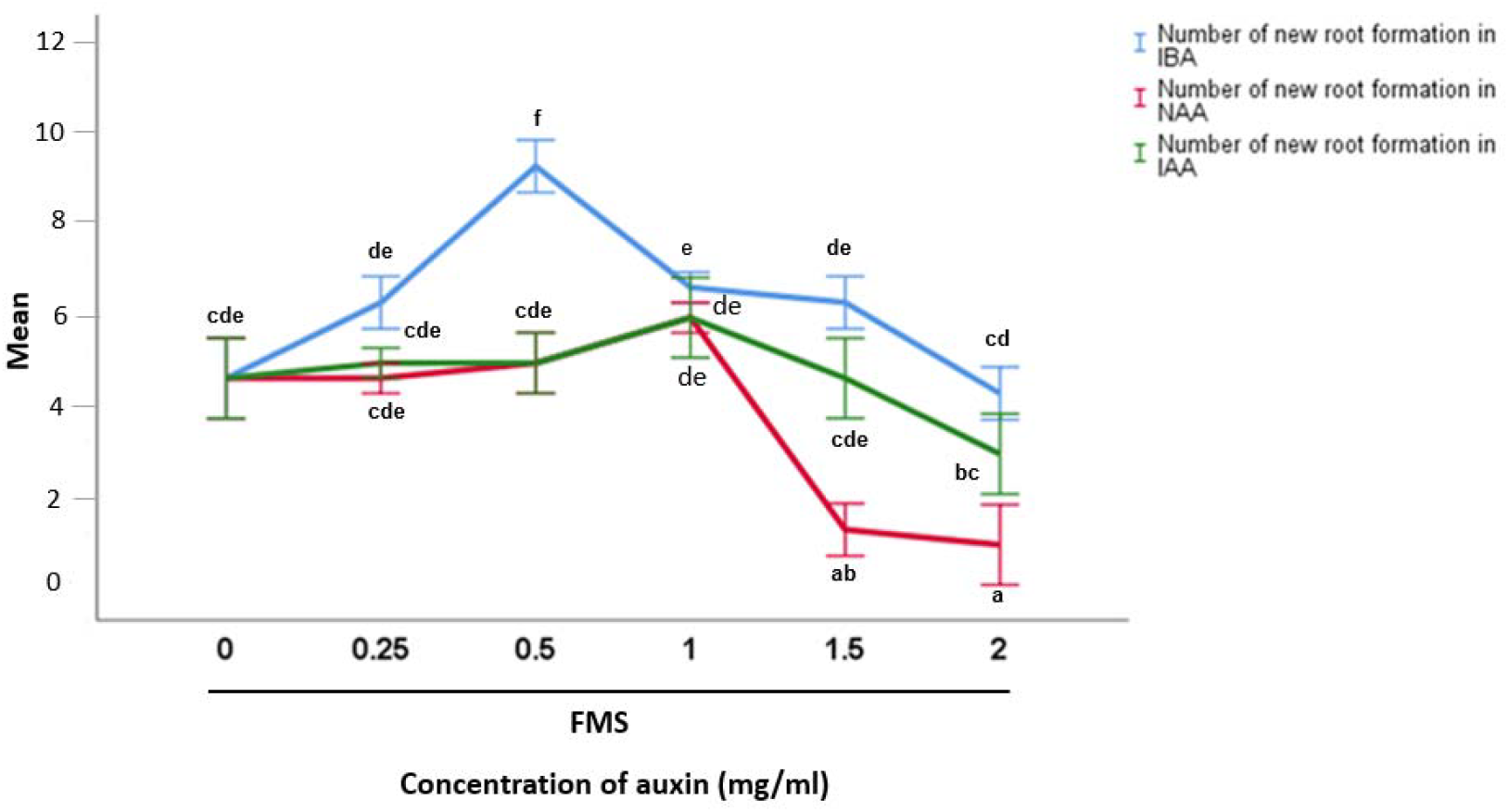
*In vitro* root development per shoot of *D. moniliforme* grown on different media combinations with different concentrations of auxin. FMS= Full murashige and skoog medium, IBA= indole-3-butyric acid, NAA= 1-naphthalene acetic acid; IAA= indole-3-acetic acid. ^a-f^ Each superscripts represents significant values at p ≤ 0.05 using Duncan’s Multiple range tests (n=36)

**Figure 4.**
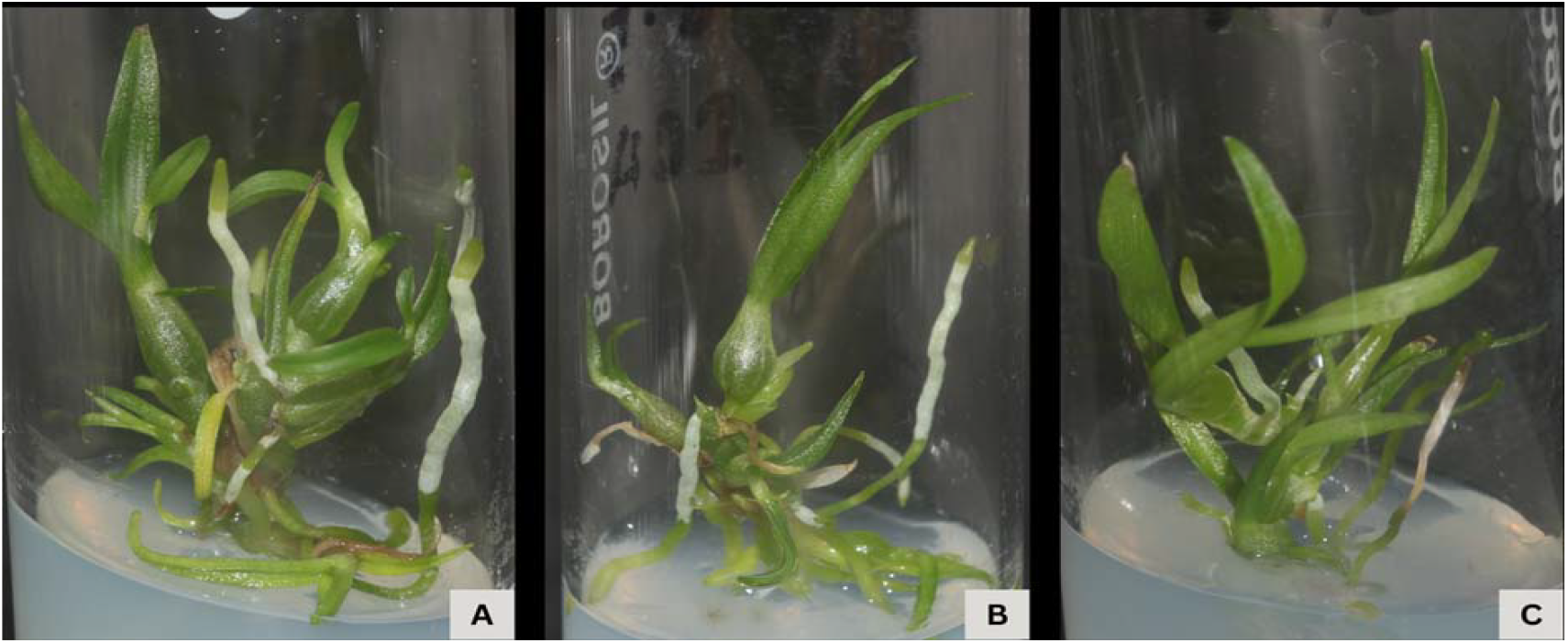
Root development of *D. moniliforme* on FMS media supplemented with different concentrations of auxin A-C: Root growth on different root induction media, **A**: FMS+0.5 mg/L IBA; **B:** FMS+1 mg/L IBA; **C:** FMS+ 0.25 mg/L IAA.

### 3.4 Acclimatization

The successful *in vivo* establishment of *in vitro*-raised Dendrobium plantlets is essential for producing high-quality commercial products, but rapid environmental changes during transfer can cause desiccation, wilting, or death without careful acclimatization (de Silva et al., 2017). To reduce mortality in the present study, healthy rooted plantlets approximately 3–5□cm in length were selected for acclimatization. After eight weeks, 70 % survival rate was found in cocopeat: moss: pine bark in the ratio□3:1:1 (Figure 5F). An important factor that influences orchid transplantation survival is the planting substrate. A good substrate has the optimum properties like water holding capacity, porosity, and drainage for the survivability of *in vitro* grown plantlets (Thapa et al., 2020). Hence this result suggests that cocopeat: moss: pinebark in the ratio□3:1:1 will be favorable for the acclimatization *Dendrobium moniliforme*.

**Figure 5.**
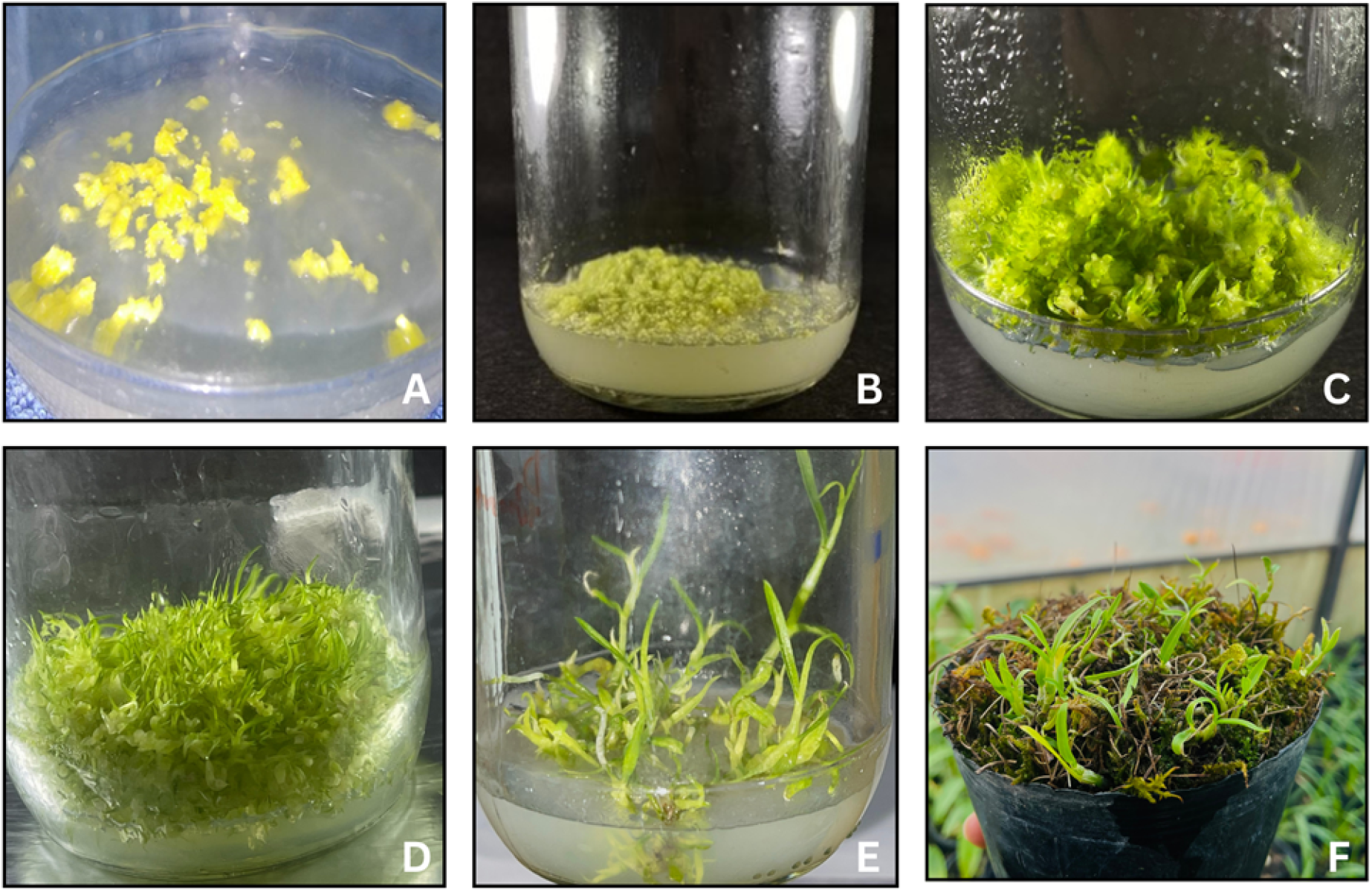
Plant regeneration of *D. moniliforme* A: inoculated seed; B: germination of *D.moniliforme*; C: Protocorm development on FMS with 10% CW; D: Shoot initiation and multiplication; E: Full grown plants of *D. moniliforme*; F: Greenhouse acclimatized plantlets of *D. moniliforme* after one month of transfer.

### 3.5 Assessment of clonal fidelity

RAPD and ISSR markers were used for the detection of genetic fidelity of the one-month successfully acclimatized plantlets of *D. moniliforme*. Template DNA from four randomly selected acclimatized plantlets and one mother *in vivo* plant was taken for the experiment. Altogether 20 Primers of the RAPD and ISSR markers were used making each of 10. Out of them, five RAPD and two ISSR markers amplified the target DNA. Monomorphic banding patterns were produced from all the amplified products. Regarding RAPD, the amplification size ranged from 200 to 1100bp. The largest amplicon size was 1100 bp on OPA-02 and OPA-03. The smallest amplicon size was on OPA-03 and OPA-06, ie. 200bp. The scorable bands produced on all the tested primers of RAPD ranged from one to five representing a total of 13 alleles. Each primer amplified an average of 2.6 alleles (Figure 6B, 6C and 6D).

**Figure 6.**
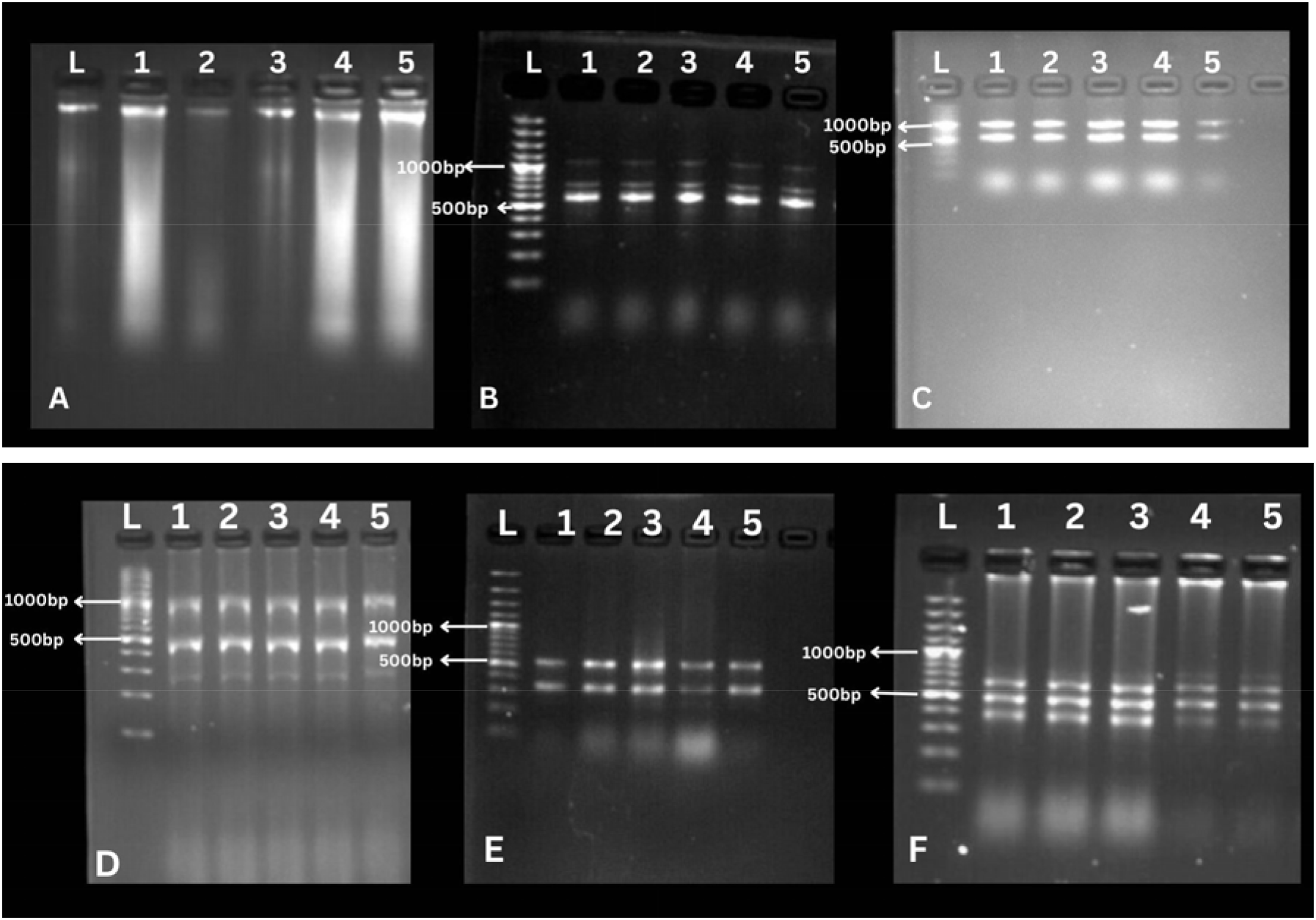
DNA profile of *D.moniliforme* on the different primers of RAPD and ISSR markers. A: DNA band; B to D: RAPD; B: OPA-02; C: OPA-07; D: OPA-03; E to F: ISSR; E: UBC-826; F: UBC: 844, L represents ladder; 1 represents amplified DNA samples of in vivo mother plant, 2, 3 & 4 represent amplified DNA samples of *in vitro* auxin and cytokinin subjected plantlets respectively; 5 represents amplified DNA samples of acclimatized plants.

Similarly, ISSR generated 5 scorable bands or alleles from genomic DNA, with amplification sizes ranging from 300 to 600 bp (Table 3). The primer UBC-844 produced the largest amplification size, whereas UBC-826 produced the smallest. The largest number of alleles (three) was amplified on UBC-844, while the minimum (two) was identified on UBC-826 (Figure. 6E, 6F). The average number of alleles amplified was 2.5 in ISSR.

**Table 3.**
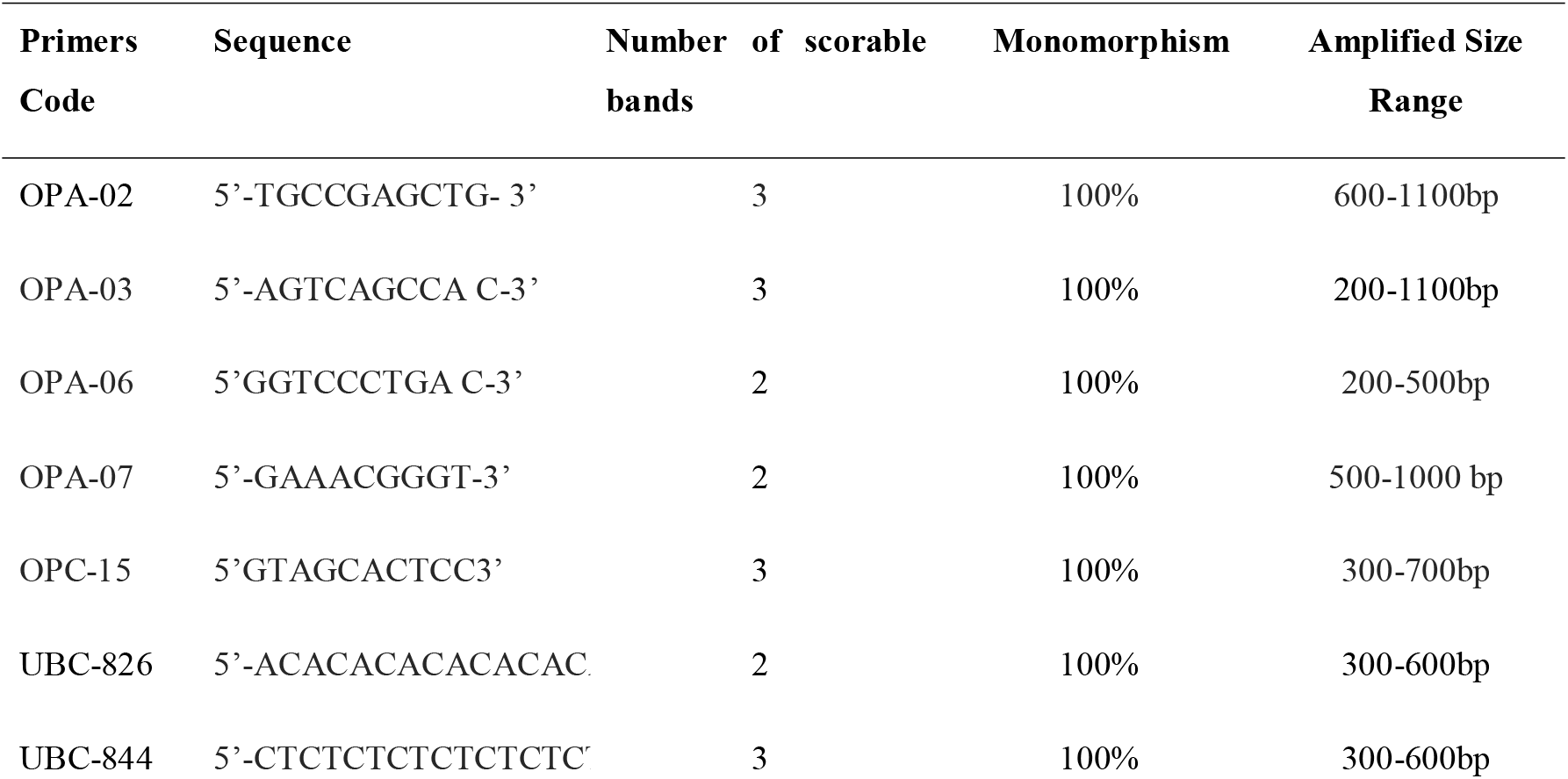
List of RAPD and ISSR primers used to confirm the genetic fidelity of *D. moniliforme*.

## 4. Discussion

Mass propagation of *D. moniliforme* for commercial usage is critical for conservation of their genetic resource. Micropropagation through plant tissue culture is a valuable alternative to conventional methods for the rapid multiplication of orchids. Seed germination and protocorm formation represent the critical and challenging steps in the orchid life cycle. Due to the absence of a functional endosperm, orchid seeds rely on a specific mycorrhizal fungus to germinate and develop (Pant et al. 2017). The orchid embryo develops into the protocorm stage upon seed germination before transforming into a plantlet. In the present study, full-strength MS media combined with 10% CW were effective for seed germination and protocorm development (five weeks and six weeks respectively). MS media was found to be stronger than HMS and QMS because it provides sufficient nutrients for early embryo development. This study aligns with Huh et al, 2016, who reported that MS medium with 10% CW improved seed germination (70.8%) and protocorm formation (74.2%) in *Cypripedium macranthos*. Previous studies have also found that coconut water is beneficial to the *in vitro* growth of various plants, including *Lindernia minima* (Tamilvanan.et al. 2022), *Dendrobium densiflorum* (Pant et al.2021), *Dendrobium transparens* (Joshi et al. 2023). Some results also suggest the addition of 15% CW for germination of *Rhynchostylis retusa* (Thomas and Michae, 2007). Coconut water acts as a natural additive which induces the growth of explants because it contains high nutritional and hormonal substances like diphenyl urea (Abbaszadeh et al. 2018).

Natural substances and hormones, like auxins and cytokinins serve as useful tools to enhance orchid developmental stages from protocorm to plantlet (Parthibhan et al. 2015). Shoot multiplication is thought to be a critical step in micropropagation where the role of phytohormones is significant (Hussain et al. 2023). The highest number of shoots developed in half-strength MS medium supplemented with 0.25 mg/l NAA and 10% CW (22±0.66 shoots/explant) and elongated shoots by 2.16cm shoots/explant. NAA helps shoots grow faster because the plant does not make enough of its auxin during regeneration (Hartati et al., 2017). Coconut water increases the possibility that organic additives could eventually take the place of plant growth regulators (Asghar et al. 2011). It is a rich source of cytokinin (Yong et al. 2009), which may favor the multiplication and differentiation of tissues in the protocorm. The present study is supported by Maharjan et al, 2020, where HMS added with CW showed a significant result for micropropagation of *D. chryseum*. According to Gnasekaran et al (2010) the addition of CW in the tissue culture is beneficial due to the presence of diphenyl urea, which is a growth factor that exhibits a cytokinin-like substance. However, some contrasting studies showed better shoot proliferation with BAP and auxins combinations, as reported by Joshi et al 2023, in *Dendrobium transparens*, Dhungana et al. 2022, in *Dendrobium crepidatum* and Pradhan et al. 2013 in *Dendrobium densiflorum*.

The rooting of *in vitro* shoots is a important step in plant micropropagation. It significantly affects the growth and survival of plantlets during the acclimatization process. In the present study, MS medium supplemented with 0.5 mg/L IBA was found to be most effective for rooting, producing an average of 10 roots/shoot. This result supports previous findings by Liu et al, 2002, who demonstrated that IBA induces better root initiation than IAA and NAA, likely due to its low toxicity and greater stability. IBA plays a major role in the processes of differentiation, elongation, and cell division (He et al. 2023). It also promotes shoot bud expansion and elongation. (Han et al. 2009; Sherif et al. 2020). Supporting our findings, other studies have also reported that IBA was found to be more effective for root development in *D. moniliforme* (Liu et al. 2023); *Vanilla planifolia* (Philip and Nainar, 1986); *Bulbophyllum odoratissimum* (Prasad et al. 2021); *Cymbidium pendulum* (Nongdam et al. 2006).

Growing orchids using tissue culture may cause some changes in the plants because of stress during the laboratory process. Clonal fidelity of plants that have regenerated should be expected since cells in culture divide mitotically during dedifferentiation and growth. But, during tissue culture, there are several factors which influence the culture condition like culture media, type of explant, sub-culture and artificial environmental conditions (Bennici et al. 2004). Another essential aspect of elite genotype preservation and its commercial implications is maintaining the genetic integrity of micro-propagated orchids. At the same time, some somaclonal variations may occur. The use of phytohormones occasionally alters the genomes of micro-propagated plants, possibly leading to point mutations, rearrangements, and DNA methylation (Bhatia et al. 2011). Thus, to confirm the quality of the plantlets for their commercial utility, it is essential to achieve genetic uniformity of micro-propagated plantlets. Clonal stability can be assessed using chromosomal counts and PCR-based molecular markers such as RAPD and ISSR (Li et al. 2019). The present study showed that micro-propagated plants of *D. moniliforme* were genetically uniform with the wild plant using RAPD and ISSR markers. It could be related to the regeneration of plantlets at lower PGR concentrations via direct organogenesis using seeds without an intermediary callus stage. Direct organogenesis produces few genetic alterations, ensuring genetic consistency (Bairu et al. 2011; Vitamvas et al. 2019; Dhungana et al. 2022). Since molecular techniques are not influenced by environmental factors, they are a more dependable and practical tool for testing the genetic uniformity of micropropagated plants than other approaches (Xiang *et al*., 2021; Huang *et al*., 2022). Sarmah et al (2024) used an RAPD marker and exhibited a uniform banding pattern in *Phalaenopsis* sp. The same result was found in Tikendra et al (2019) used 15 RAPD and 11 ISSR markers to check the genetic uniformity of *D. chrysotoxum*. They got 100% monomorphism with the mother plant. Similiarly, using RAPD and ISSR markers, for some medicinal plants, such as *Tinospora cordifolia* (Pandey et al. 2023) and Piper Longum (Thapa et al. 2024), it was found that each of the *in vitro* regenerated plantlets were genetically identical to the mother plants. Basyal et al 2025, used RAPD and ISSR to evaluate the genetic integrity of in vitro regenerated *Curcuma aeruginosa* plantlets and found 100% monomorphism with the mother plant.

## 5. Conclusion

Micropropagation of *Dendrobium moniliforme* (L.) Sw. was found to be related to various phytohormones and natural additives. MS media supplemented with 10% CW was crucial for seed germination and protocorm formation. Whereas Half MS media supplemented with NAA and coconut water was required for *in vitro* shoot development, IBA was required for rooting during micropropagation of *D. moniliforme*. Furthermore, the regenerated plantlets were genetically uniform with the mother plant based on molecular markers. This optimized protocol for the mass propagation of *D. moniliforme* is wise with sustainable production, as well as, germplasm conservation. This result serves as a starting point for further research into the micropropagation of endangered orchids.

## Author’s contribution

Manisha Ghimire: Investigation, writing-original draft, data curation, Mahendra Thapa: Methodology, data curation and investigation. Mukti Ram Paudel, Pusp Raj Joshi, Sven H. Wagner, Pritam Gurung, Krishna Kumar Pant: Review and editing, formal analysis, Bijaya Pant: Writing, review and editing, supervision, and conceptualization.

## Acknowledgement

The authors are thankful towards the Central Department of Botany, Tribhuvan University and Annapurna Research Center for providing space for laboratory work. The authors are pleased to acknowledge Puskar Basyal, Lasta Maharjan and Sibesh Maharjan for helping me throughout the research.

---

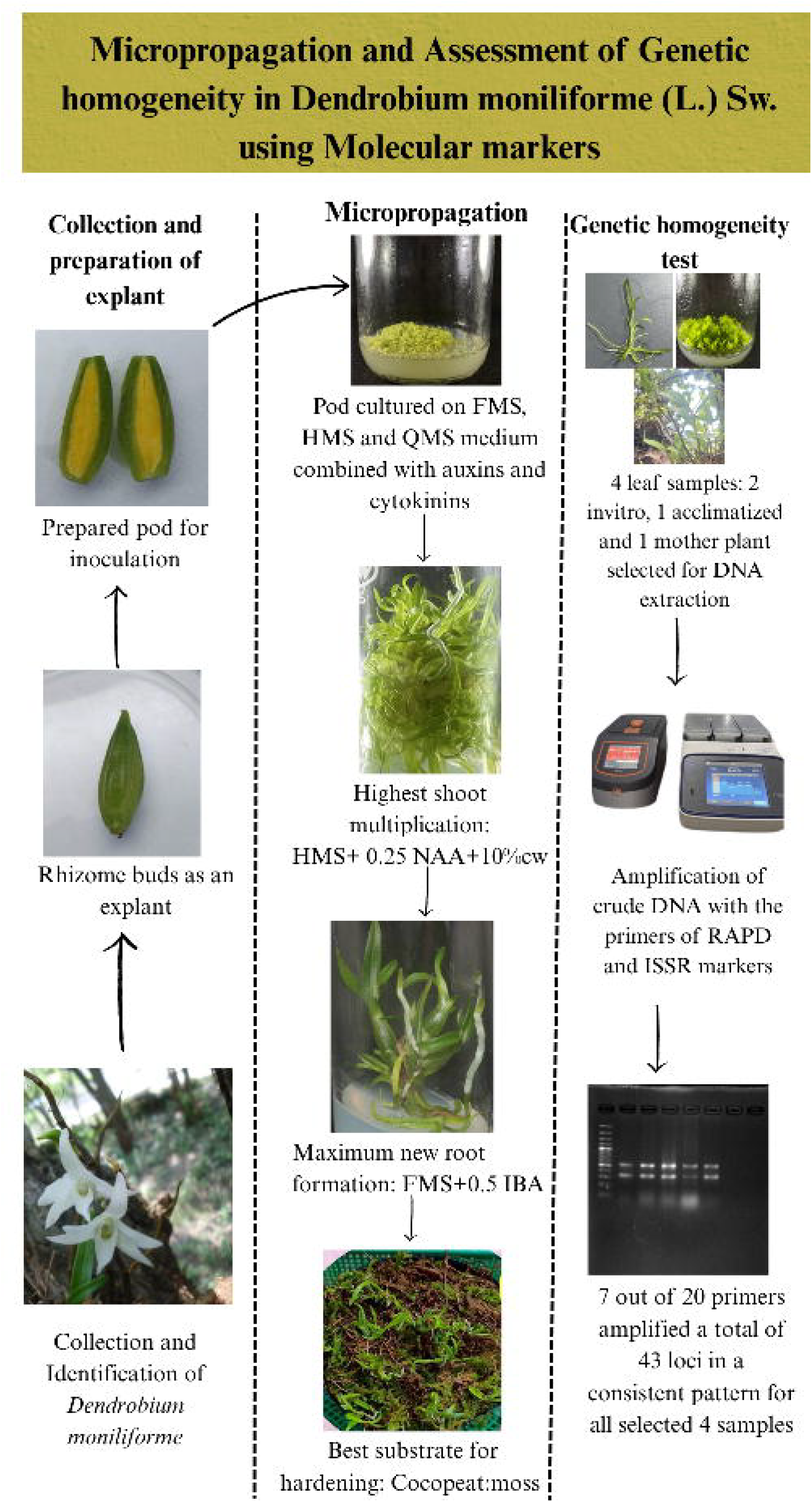

